# Homologous chromosome recognition via nonspecific interactions

**DOI:** 10.1101/2023.06.09.544427

**Authors:** Wallace F. Marshall, Jennifer C. Fung

## Abstract

In many organisms, most notably *Drosophila*, homologous chromosomes in somatic cells associate with each other, a phenomenon known as somatic homolog pairing. Unlike in meiosis, where homology is read out at the level of DNA sequence complementarity, somatic homolog pairing takes place without double strand breaks or strand invasion, thus requiring some other mechanism for homologs to recognize each other. Several studies have suggested a “specific button” model, in which a series of distinct regions in the genome, known as buttons, can associate with each other, presumably mediated by different proteins that bind to these different regions. Here we consider an alternative model, which we term the “button barcode” model, in which there is only one type of recognition site or adhesion button, present in many copies in the genome, each of which can associate with any of the others with equal affinity. An important component of this model is that the buttons are non-uniformly distributed, such that alignment of a chromosome with its correct homolog, compared with a non-homolog, is energetically favored; since to achieve nonhomologous alignment, chromosomes would be required to mechanically deform in order to bring their buttons into mutual register. We investigated several types of barcodes and examined their effect on pairing fidelity. We found that high fidelity homolog recognition can be achieved by arranging chromosome pairing buttons according to an actual industrial barcode used for warehouse sorting. By simulating randomly generated non-uniform button distributions, many highly effective button barcodes can be easily found, some of which achieve virtually perfect pairing fidelity. This model is consistent with existing literature on the effect of translocations of different sizes on homolog pairing. We conclude that a button barcode model can attain highly specific homolog recognition, comparable to that seen in actual cells undergoing somatic homolog pairing, without the need for specific interactions. This model may have implications for how meiotic pairing is achieved.

## Introduction

In meiosis, homologous chromosomes are thought to recognize each other at the level of DNA sequence. Specialized enzymes create double strand breaks, from which single strands extend to test homology with other chromosomes. This process of sequence-based homology assessment leads to a highly precise alignment of each chromosome with its correct homolog, allowing for recombination between homologous loci to establish crossovers for proper segregation during meiosis I division.

However, in many other situations outside of meiosis, homologous chromosomes can recognize and associate with each other, despite a lack of double strand breaks, single strand invasion, or nucleotide-level base pairing. The association of homologous chromosomes in non-meiotic cells is known as somatic homolog pairing, and it has been reported in many different organisms and cell types including in humans (Stevens 1908; Metz 1916; Arnoldus 1989; Apte 2012; Joyce 2016). In some cases, only small chromosomal segments associate with their homologs, and this can vary between cell types or disease states. Somatic pairing is most extreme in dipterans such as *Drosophila*, in which homologous chromosomes are paired in virtually all tissues after the first 13 cell cycles of early embryos (Stevens 1908; Metz 1916; Hiraoka 1993; Fung 1998).

The physiological purpose of somatic homolog pairing is unknown. In cases where recombination independent association takes place prior to meiosis, it may be involved in setting the stage for a more precise alignment once double strand breaks have formed. Somatic homolog association might also facilitate DNA repair by homologous recombination in G1 when sisters are not yet available for this purpose, by placing homologs near each other. In some instances, physical association of chromosomes is involved in trans-regulation of gene expression by regulatory elements located on the other chromosome (Fukaya 2017; Joyce 2019), and pairing may correlate with chromosome functional state (AlHaj Abed 2019).

Regardless of the physiological function of recombination-independent association, an equally interesting question is its mechanism. Genetic analyses of transvection and pairing in flies carrying translocations and other chromosome rearrangements have shown that large chromosome regions, rather than specific DNA elements such as enhancers or insulators, are involved in assessing homology (Lewis 1954; Ou 2009; Viets 2019). This has led to the idea of a “Specific Button” model, in which chromosome regions sparsely distributed along chromosome arms mediate specific associations (Viets 2019). This “Specific Button” model is consistent with both FISH (Fung 1998) and live-cell imaging (Child 2021) studies of somatic pairing in *Drosophila*, which showed that chromosomes do not “zip up” continuously along their length, but instead initiate pairing independently at multiple distinct regions.

In a recent tour-de-force study of the kinetics of somatic pairing in *Drosophila*, Child and co-workers (2021) implemented a computational version of the button model for pairing, in which a set of discrete pairing sites distributed at regular intervals along the chromosome could engage in independent pairing interactions that, collectively, would align the two homologs along their length. Computational modeling indicated that this model can account for pairing kinetics consistent with live cell rate measurements, but it only works if the buttons are distinct, in the sense that a button at a given position on a chromosome can only pair with a corresponding button on the homologous chromosome. A model in which the buttons lacked specificity was not able to achieve homologous recognition (Child 2021), although it has been shown that regularly spaced nonspecific association sites are able to at least bring chromosomes into alignment, albeit without any specificity (Nicodemi 2008). The lack of specificity is simply because there is no energetic difference between associating with the correct versus incorrect pairing partner (**Figure 1**).

Inspired by the work of Viets (2019) and Child (2021), we investigated a variant of the button model, in which the pairing “buttons” are individually non-specific, such that every button is equally capable of pairing with any other button, but in which the buttons are non-uniformly arranged in a different pattern on different chromosomes, allowing them to act as a code specifying chromosome identity (**Figure 1**). We will refer to this model as the “Button Barcode” model. Specificity of homolog pairing would require the chromosome pairing process to be able to distinguish the spacing of these non-specific buttons over a potentially long spatial scale. We propose that the physics of the chromatin polymer will tend to favor association of buttons that are equally spaced on both chromosomes. Association of pairs of buttons with different spacings on two chromosomes will require one or both chromosomes to either stretch or compact, incurring a mechanical energetic cost. In such a model, the information about chromosome identity is encoded in the spacing between buttons, in much the way that an industrial barcode encodes information in the spacing between bars printed on a package (Palmer 1995). The key feature of this model is that the spacing between the buttons, not the position of buttons per se, is the origin of selective association.

Here we use a coarse-grained computational model of chromatin as a worm-like chain to investigate the plausibility of this button barcode model. Our model is not intended to represent any particular species or model system, but just to reflect generic properties of chromosome polymers. We show that unequal spacing of non-specific interaction sites on different chromosomes is in fact sufficient to produce a reliable association of chromosomes with their homologous partners. We show that this specificity depends on the mechanics of the chromosomes; on the three dimensional organization of chromosomes within the nucleus, specifically the Rabl configuration in which centromeres cluster at one end of the nucleus and telomeres at the other; and on the reversibility of pairing interactions. We show that randomly spaced buttons are able to achieve a level of specificity that matches what is seen in actual cases of somatic pairing. Finally, we implement a chromosomal version of a standard industrial barcode known as “code 2 of 5” (Allais 1984) and show that it out-performs many random button patterns. Our results show that, at least in principle, sequence-level specificity is not required for accurate homologous pairing, and suggest some features that would be required for this type of mechanism to work. We discuss this model in light of evidence for and against specific buttons, and conclude with a model in which barcode segments built from small non-specific interaction buttons can act like specific buttons at a larger scale, thus potentially explaining the existing data concerning the effect of translocations on pairing while avoiding the need to posit specific interactions at a molecular level.

## Methods

### Coarse grained polymer model for chromosome dynamics

The model used is based on a previous coarse-grained Langevin model developed for meiotic chromosome movement (Marshall and Fung 2016; Marshall and Fung 2019). In this model, each chromosome is represented as a string of beads joined by springs. Each bead (node) is subject to a random thermal (Langevin) force, a frictional force proportional to velocity, and additional force terms that describe the physical properties of the chromosome and its interaction with the nuclear envelope.

These forces yield the following equation of motion, which is used to update the velocity and position of each node at each time step:

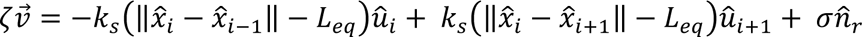

where x_i_ and v_i_ are position and velocity vectors corresponding to node i on the chain, ζ is the friction coefficient, *k*_s_ and *L*_eq_ are the spring constant and equilibrium length of the links, σ is the average magnitude of the Langevin (thermal) random force, and n_r_ is a randomly oriented unit vector generated for each node at every time point, and u_i_ represents a unit vector directed from node i-1 to node i. We represent a worm like chain using the Kratky-Porod model in which a force is generated proportional to the bending angle between two successive links with a proportionality constant k_bend_.

The Langevin random force actually represents the average net resultant of all random collisions with thermally excited solvent molecules during a single time-step. The magnitude of the Langevin random force is assumed to be constant at every time step, so σ is constant. At each time point, a new orientation is chosen for each of the random unit vectors n_r_ is chosen from a uniform distribution of angles in spherical coordinates.

In this model, we employ a Rouse “phantom polymer” model in which the chromatin chains are able to freely pass through each other. This assumption is supported by polymer scaling arguments based on the relative density of topoisomerase strand-passage sites compared to the density of chromatin interlocks, which indicate that any interlocks that form would be rapidly resolved by topoisomerase activity (Sikorav and Jannink, 1994), a behavior that is also supported by experimental measurements of topoisomerase ability to convert an entangled DNA melt into a viscous fluid (Kundukad and van der Maarel, 2010). While this assumption may not be appropriate for meiotic chromosomes, which consist of four strands each and in many cases also contain protein-based axial elements, which would prevent strand passage and therefore require a model that can capture entanglement effects (Navarro 2022), in the case of somatic homolog pairing, these concerns do not apply and so the Rouse assumption is justified. During motion, the chromosomes are confined inside a spherical nuclear envelope by a repulsive force applied to any node moving outside the spherical shell in a direction normal to the surface with a spring constant k_nuc as previously described (Marshall and Fung, 2016).

To simulate chromosome dynamics during pairing, a chain is initialized to a random configuration using a lattice model, and then relaxed according to the Langevin equation and nuclear constraints, as previously described (Marshall and Fung, 2016). After the initial relaxation period is over, subsequent timesteps execute the following steps. First, a test is made to see if any of the currently paired nodes will become unpaired. Then, the displacements calculated for each node in the previous iteration are applied to all nodes. After the node positions are updated, a test is made to see if any nodes in the pairing list are within a capture radius of each other, and if so, they are assigned to the paired state with probability p_pair. Finally, new velocities are calculated for every node, taking into account the current set of node positions, according to the equation above.

### Modeling nonspecific interactions

For each simulation, a list of nodes capable of pairing (the “buttons”) is specified. As the bead-spring chains undergo their random movement, whenever two loci on the button lists for any of the chromosomes move to with a defined capture radius of each other, they are set to a paired state. This pairing process does not discriminate between buttons on different chromosomes - any button is allowed to pair with any other button, including on the same chromosome, its correct homolog, or either of the other chromosomes. When a button is set to the paired state, the identifier of the button to which it is paired is stored, and from then on the motion of the two nodes will subsequently be forced to coincide (details for enforcing this correlated motion are given in previous work, Marshall and Fung 2016). Once a pair of buttons are set to the paired state and assigned to each other, they are no longer included in subsequent pairing tests.

At subsequent time-steps, unpairing of paired loci occurs with a fixed probability given by the unpairing probability p_unpair. In this model, the value of p_unpair is the same for all pairs of buttons, so that discrimination between correct and incorrect associations can only be achieved by higher order structural or mechanical influences, and not on the local interaction kinetics of the buttons themselves. If two nodes become unpaired, they are set to the unpaired state but not moved apart. The unpairing test is done at the start of each iteration, and whether the nodes will rapidly re-pair depends on whether their random motions during the timestep take them further apart than r_capture during the iteration.

### Modeling the Rabl configuration

In many cells, chromosomes are arranged in a Rabl configuration, in which telomeres cluster together at one end of the nucleus and centromeres at the other (Comings 1980). This is conspicuously true in *Drosophil*a embryos at the time of establishment of somatic pairing (Marshall 1996; Dernburg 1996). The Rabl configuration arises as a remnant of chromosome organization during the preceding anaphase, in which the centromeres move together to each pole of the spindle. In addition, chromosomes are often associated with the nuclear envelope (Brickner 2017), with specific chromosome regions, such as telomeres or heterochromatic repeats, non-randomly located in the nuclear periphery. Again this is particularly evident in *Drosophila* embryos (Marshall 1996). We represent the Rabl configuration by confining the centromeres, defined as the midpoint of each chromosome polymer, to a restricted region on the nuclear surface by applying a restoring force, proportional to the distance to the boundary of the region, to any centromeres that move outside of the region. The size of this region is given by the parameter confinement_radius. Within the region, the centromeres are subject to the standard Langevin random force of the model, but they are prevented from leaving the region of constraint.

### Parameter choice

The model presented here is intended to ask whether non-specific interactions can, in principle, provide specificity of homologous association. It is not intended to accurately represent the details of any specific biological system. Here we present the choices for parameter values used in this model, in which we have tried to assume simple values that are consistent to a first order with biologically measured values, when such are known.

Within the model, we need to define units of length, time, and force. We begin by determining units that make our assumed values of link length, time step, and Langevin force have reasonable values. We then use these units to convert our other parameters into real units, which we can then compare with estimates in real systems.

#### Link length (L_eq_) 1 200 nm

To estimate the link length in the bead-spring chain, we note that each arm of the chromosome has 25 links. We previously measured the position of multiple loci on one arm of *Chromosome* 2 in *Drosophila* (during cycle 13) and found that the chromosome spans a linear distance of 4 microns (Marshall 1996), which would correspond to 160 nm per link. For our simplified model, we assume a slightly greater link length of 200 nm. Given that a *Drosophila* chromosome is approximately 50 Mb in size, each 200 nm link would correspond to approximately 1Mb of DNA. Sun (2021) has reported that the size range of topological associated domains (TADs) in *Drosophila* embryos is in the range of 0.3-1 Mb, roughly consistent with our model assumptions.

#### Time Step 1 10 ms

To estimate what length of time each step of the simulation corresponds to, we start by noting that the total simulation is carried out for 300,000 steps. The duration of cycle 14 in *Drosophila*, when somatic pairing occurs, is approximately 1 hour. 300,000 steps per hour corresponds to 12 ms per step. We simplify here by setting the time step to 10 ms.

#### Langevin force 0.15 0.3 pN

To estimate the Langevin force used in our simulation, we start with the measured value of D=2.0×10^−11^ cm^2^/sec for the diffusion coefficient of interphase chromatin measured experimentally in *Drosophila* embryos (Marshall 1997), which is equal to D=2000 nm^2^/sec. We then use the relation δ=sqrt(6ΔtD), to get the root mean-squared (rms) displacement during a single 10 ms time step:

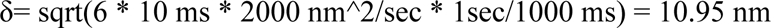

which we approximate as 11 nm. We make the assumption that during a single time step, this displacement is driven by continuous application of thermal energy of magnitude k_B_T, which we take as approximately 4 pN nm at 30C.

Thus, the Langevin force is k_B_T/δ = 4 pNnm / 11 nm = 0.26 pN, which we approximate as 0.3 pN.

#### Unit Conversion Factors

With these three values estimated, we can compare the parameter values used in the simulation to the physical values they correspond to, and thereby compute the units of all other parameters in the simulation. For the force units, we divide the actual Langevin force (0.3 pN) by the Langevin force parameter used in the simulation (0.15 force units). The unit conversion factors are then as follows:

1 time unit = 10 ms
1 distance unit = 200 nm
1 force unit = 2 pN

#### Friction coefficient (σ)1.0 3×10^−4^ pN s /nm

To derive the frictional coefficient ζ, we note that a node of the chain moves by δ=11 nm in a single timestep corresponding to 0.01 s, creating an apparent velocity on the order of 1000 nm/s. This motion is assumed to be driven by the Langevin force of 0.3 pN acting continuously during the time step, driving the node at a constant velocity. From the relation ζ=σ/v we obtain a value for ζ of 0.3 pN / (1000 nm/s) = 3×10^−4^ pN s/nm which is equivalent to 3×10^−7^ Kg/s.

To see if this is reasonable, we can calculate the viscosity corresponding to this frictional coefficient assuming Stoke’s law for a perfect sphere, such that η= ζ/6πR. Assuming the de Gennes model of spherical globules whose diameter matches the link length (Grosberg and Khoklov 1994), we take R=100 nm = 10^−7^ m, which yields a value for viscosity of (3×10^−7^ Kg/s)/(6*3.14*10^−7^ m) = 0.08 Kg/ms = 4 cP.

We thus estimate a viscosity that is on the order of four times more viscous than water (viscosity 1 cP), consistent with prior studies of the viscosity for the fluid phase of nucleoplasm (Peters 1984; Erdel 2015; Speil 2010).

#### Spring constant (k_s_) 0.25 0.0025 pN/nm

The parameter k_s_ is a spring constant describing the force needed to extend a single link in the chain by 1 length unit. To put this into real units we take the product 0.25 force units / length unit * 2pN/force unit / (200 nm/length unit) = 0.0025 pN/nm.

#### Torsion spring constant (k_bend) 0-0.4 0 - 0.8 pN

Our wormlike chain model imposes a force component to oppose bending of the chain. We chose the range of values for the bending parameter k_bend empirically to generate a relevant range of persistence lengths, via simulations with a large (100 length units) nuclear radius to avoid interactions with the NE. From these simulations, we observed a linear relation, with Lp = 40*kbend.

Our minimum value for k_bend is zero, in which case no penalty is imposed for bending the chain, and the model becomes a freely jointed chain. In this case, the persistence length Lp is 100 nm. Our maximum value of kbend is 0.4 force units, which corresponds to 0.8 pN for k_bend.

#### Nuclear Envelope Spring Constant (k_nuc) 0.35 3.5 pN per micron

We represent the nuclear boundary as an elastic shell such that any node of a chromosome chain that moves beyond the specified nuclear radius experiences an inwardly directed radial force proportional to the distance it has moved past the radius. Loci located at or within the nuclear radius do not experience any such force. Representation of the NE as a linear spring is consistent with prior mechanical measurements of nuclear deformation (Dahl 2004; Kaufmann 2011; Stephens 2017; Zuela-Sopilniak 2020). Measurements using AFM and optical tweezers to probe the apparent spring constant of the nuclear surface in Xenopus oocytes (Kaufmann 2011) and mouse embryonic fibroblasts (Vahabikashi 2022) have yielded values in the range 5-20 pN/micron.

#### Nuclear radius (nuc_radius) 15.0 3 microns

The nucleus radius of 15 length units corresponds to 3 microns, comparable to the size scale of cycle 13 nuclei. During cycle 14, nuclei become narrower but lose their spherical shape and become elongated. Here, for simplicity, we assume a spherical nuclear shape.

#### Unpairing probability (P_unpair) 0.4

P_unpair 0.2 means that the average time two loci spend in the paired state, after pairing would be 5 time units, which is just 50 ms. This is orders of magnitude shorter than the reported times that paired loci spend in the paired state (Lim 2018; Child 2021). However, the avidity effect means that most of the time, a pair of buttons that became unpaired would re-pair so rapidly that the event would not have been detected in live imaging. Unless the loci paired were the only paired loci on a chromosome, they would be highly likely to re-associate rather than drift apart. Standard methods for visualizing chromosome unpairing, based on the presence of one versus two distinct spots in fluorescence imaging, would not in general be able to detect short transient unpairing events. As a technical note, the check for unpairing is done at the start of each simulation time-step, after which the nodes are displaced according to the forces previously calculated for each node. If the displacements are small, the nodes may still be within the capture radius of each other, and thus be able to re-pair with probability p_pair before the forces are calculated for the next iteration.

#### Pairing probability (P_pair) 1.0

We assume that whenever two buttons come within the capture radius of each other, they will pair with a defined probability. As noted above, this pairing is reversible. Once two nodes are paired, their subsequent motion is forced to be equal, with forces on the pair of nodes obtained as the resultant of the forces applied to each of the two nodes in the pair.

#### Capture distance (Dist_capture) 0.5 100 nm

The capture distance was chosen to allow sites to pair when the polymer globules represented by the links are overlapping by the radius of the globule. Under the standard De Gennes globule model, the diameter of a globule equals the link length of a bead-spring representation, so we take half a link length as the capture radius.

#### Confinement radius 2.0 400 nm

This parameter reflects the size of the region of confinement of the centromeres which is used to model the Rabl configuration. In *Drosophila* embryos, centric heterochromatin of all the chromosomes is localized in “chromocenters” that are on the order of 1 micron in diameter (Jagannathan 2018; Jagannathan 2019). Unless noted otherwise, all of the simulations reported here used a small confinement radius of 2.0 length units corresponding to 400 nm. However, in simulations adjusting the confinement radius (Figure 3C), no real effect was seen on the Rabl correlation coefficient, our metric for the strength of Rabl alignment of the chromosomes along the z axis, until the radius was greater than 10 length units, corresponding to 4 microns. Thus, the size of the confinement region required to maintain an effective Rabl orientation is well within the range of sizes consistent with the size of *Drosophila* chromocenters.

#### n_steps 300000 50 min

All simulations were run for a total of 300,000 timesteps. Given that each step represents 10 ms, this is equivalent to 50 minutes of real time. The entire duration of cycle 14 in the *Drosophila* embryo is roughly 1 hour, so this is comparable to the time normally available to achieve a high level of somatic pairing.

#### initial_relaxation 20000 200 s

At the start of the simulation, each chromosome is initialized by picking a random point inside the nucleus as the first node, and then adding additional nodes according to a random walk, with the choice of node positions limited to points inside the nucleus. Once this initial configuration is generated, the simulation is run for 20000 iterations, without any pairing, in order to allow the chromosome to relax into a configuration that is consistent with the simulated forces. This number of steps was chosen based on simulations in which the nuclear envelope repulsion force was turned off and the simulation run until the end-to-end distance distribution had reached a steady state (Marshall and Fung 2016). After the initial relaxation phase, the simulation continues to run but now pairing is turned on.

### Random button code generation

To generate random button patterns, we used the randperm() function of Matlab to permute a list of n nodes, and then took the first m to be buttons which were then input to the simulation program. In the case of random codes for comparison to the 2 of 5 code, for which there is always a button at both ends of the region in question, we assigned the endpoints of the region to be buttons, and then applied the random permutation scheme to the intervening n-2 nodes.

### Implementing an industrial barcode

To convert code 2 of 5 (Allais 1984) into a button pattern spanning a chromosome arm, we assign a button to node 25 (the centromere) and then add additional buttons at spacings given by the following rule: for a narrow bar in 2 of 5, the next button is spaced 2 nodes away. For a wide bar in 2 of 5, the next button is spaced 6 nodes away. This rule preserves the 1:3 ratio of narrow to wide bars used in the industrial code. The resulting button patterns corresponding to the digits in 2 of 5 are as follows, where the binary strings represent the normal way that 2 of 5 is written in text, with 0 corresponding to a narrow bar and 1 to a wide bar (Harmon and Adams 1989)

|  |  |  |
| --- | --- | --- |
| 0: | 00110 | 25, 27, 29, 35, 41, 43 |
| 1: | 10001 | 25, 31, 33, 35, 37, 43 |
| 2: | 01001 | 25, 27, 33, 35, 37, 43 |
| 3: | 11000 | 25, 31, 37, 39, 41, 43 |
| 4: | 00101 | 25, 27, 29, 35, 37, 43 |
| 5: | 10100 | 25, 31, 33, 39, 41, 43 |
| 6: | 01100 | 25, 27, 33, 39, 41, 43 |
| 7: | 00011 | 25, 27, 29, 31, 37, 43 |
| 8: | 10010 | 25, 31, 33, 35, 41, 43 |
| 9: | 01010 | 25, 27, 33, 35, 41, 43 |

The simulations in Figure 6 used code 2 of 5 symbols for 0 and 1.

## Results

### Modeling somatic chromosome interactions

The essence of the Button Bar Code model is described in **Figure 1B**. In this model, any button can pair with any button, and selectivity for the correct homolog is a consequence of the different arrangement of the buttons along the different chromosomes combined with the physics of the chromatin polymer. Although chromatin is sometimes modeled as a freely jointed random chain, actual chromosomes are in many cases better described as worm-like chains or other forms of elastic polymers, in which energy is required to bend them away from an equilibrium (Ehrlich 1997; Marko and Siggia 1997; Bystricky 2004; Penfold 2012). The end-points of any segment of such a worm-like chain will have a characteristic distribution of lengths that is energetically favorable. Trying to move the ends of that segment closer together or farther apart will incur an energetic cost. Because of this energic cost to deforming (either looping or stretching) the polymer, a side-by-side alignment of actual homologs should be energetically favored over alignment of non-homologs because only an association of actual homologs allows the segments between adjacent pairs of buttons to remain at their energetically favored lengths. If non-homologs attempt to associate, it would require a bending or stretching of one or both homologs in order to accommodate the mismatch in spacing between adjacent buttons.

**Figure 1.**
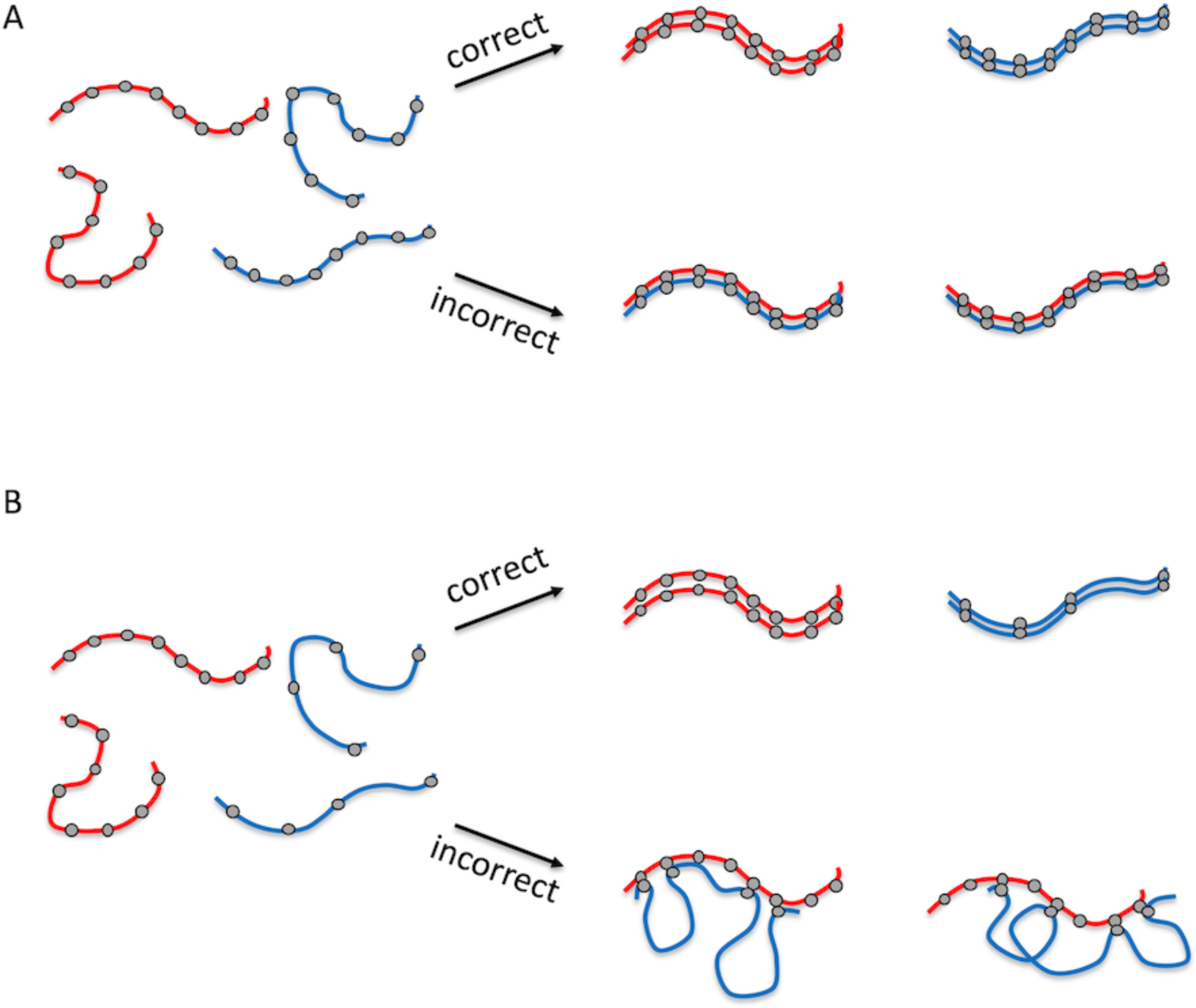
A button barcode model for homolog recognition by non-specific interactions. **A.** Uniformly spaced non-specific pairing buttons. In this case, grey circles denote pairing buttons, each of which has the same molecular affinity for every other such button in the genome. Two different pairs of homologs are denoted by red and blue color. Correct pairing (red pairs with red, blue pairs with blue) and incorrect pairing (red pairs with blue) would each be equally likely. **B.** Button bar-code model, in which non-specific pairing buttons have different spacing patterns on the two chromosomes. In this case, pairing with the incorrect homolog incurs an energetic cost for deforming one or both chromosome polymers so as to allow the buttons to physically interact. Pairing with the correct homolog does not require such deformation and is thus energetically favored, hence more probable.

In order to test the plausibility of this model, we implement a Brownian dynamics simulation based on prior modeling of meiotic chromosome movement and pairing (Marshall and Fung 2016; Marshall and Fung 2019). As illustrated in **Figure 2A**, we represent the chromosome using a bead-spring model, with each node (bead) subject to a Langevin random force that represents thermal energy, as well as forces applied by the springs linking that node to its two neighbors. We also impose a series of torsion spring forces that tend to push the chromosome towards an elongated linear form, creating a worm-like chain model (**Figure 2B**). Finally, the bead-spring chains are confined to a spherical nucleus, within which centromeres are clustered in a limited region of the nuclear surface to mimic the Rabl configuration (**Figure 2C**).

**Figure 2.**
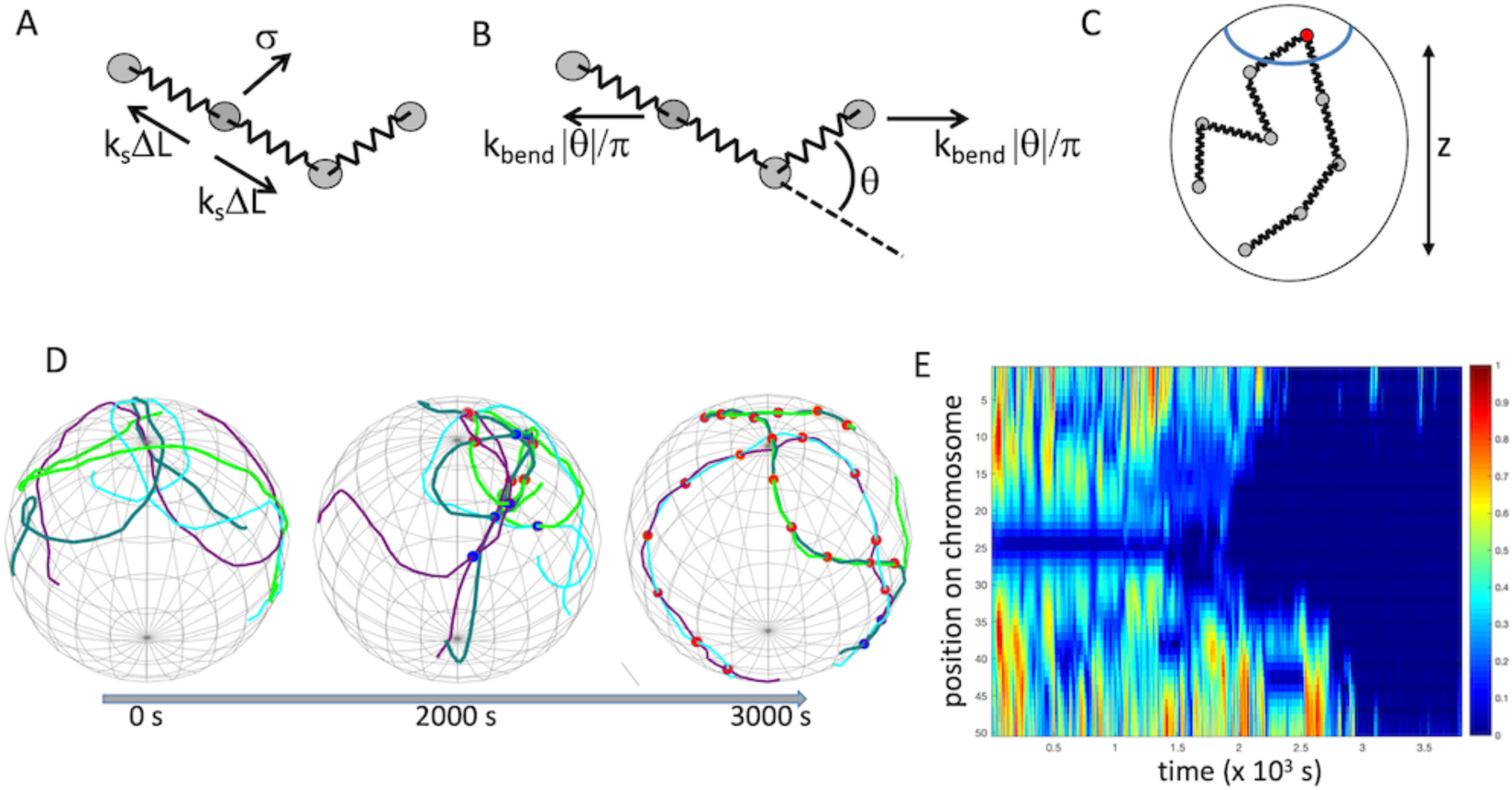
Computational model for somatic homolog pairing. **A.** bead-spring model for chromosome polymer dynamics. Chromosome is represented by beads linked by Hookean springs with spring constant k_s_. Each bead, which we term a node, is subject to a random Langevin force 0, as well as forces generated by the stretching of the springs linking it to adjacent nodes. The entire chain is modeled as moving in a medium with a specified frictional coefficient. **B.** Chromosome flexibility is modeled using a worm-like chain model in which adjacent nodes are pulled apart by a force proportional to the deviation of the chain shape from a straight line. The higher the bending constant, the less flexible the chain and the more it tends to adopt an extended shape. **C.** Chromosomes are modeled as confined within a spherical nucleus. The centromere of each chromosome, indicated by the red node, is held attached to the nuclear envelope within a confined surface patch denoted by the blue circle. This centromere clustering creates a Rabl orientation and defines a vertical axis for the nucleus, which we denote as the z axis in our simulations. **D.** Image sequence from a representative simulation in which buttons are present at a regular spacing on the two chromosomes, but with different spaces on the two chromosomes, with buttons located every three nodes on one chromosome and every four nodes on the other, spanning the entire chromosome in each case. Two pairs of homologs were simulated, with one pair of homologs plotted in dark and light green, and the other in cyan and purple. Incorrectly paired loci are marked with a blue sphere, and correctly paired loci are marked with a red sphere. **E**. Pairing kymograph from the same simulation as panel D, plotting distance between homologous loci for each position along one chromosome. X and Y axes denote timesteps of simulation and position along chromosome, respectively. The color code uses a jet colormap to map the distance of each locus to its homolog, normalized to the maximum distance between all homologs observed in the simulation, with red indicating maximum distance, and dark blue representing zero distance (which corresponds to the paired state). The color bar on the right gives the colors as a function of fraction of maximum distance.

**Figure 2D** shows three time points in a representative simulation, in which the buttons are spaced at regular intervals on both chromosomes, but with different spacings (three nodes apart on one chromosome, four nodes apart on the other). Early in the simulation, chromosomes have not yet paired. Later, buttons have begun to associate with each other, some with the correct (homologous) button, but others with incorrect buttons. This is not surprising given that there is no selectivity in terms of which buttons are allowed to pair with other buttons. As the simulation proceeds, fewer and fewer incorrect associations are seen, and more and more correct associations are seen. A pairing kymograph plot (Marshall and Fung 2016), which depicts the distance of each locus to its homolog over time, shows that a set of completely non-specific buttons, arranged with unequal spacing on the two chromosome pairs, ends up producing close spacing of all loci with their homologous loci. (**Figure 2E**).

We note that while our model is roughly based on observations of somatic homolog pairing in *Drosophila* embryos, the model itself is highly simplified and is not meant to represent any particular species or cell type. Instead, the goal is just to test the plausibility of such a model and explore what features a chromosome would need to have in order to achieve a sufficiently high degree of correct homologous pairing.

### Influence of polymer mechanics on pairing specificity

We next investigated the influence of several key model parameters on the ability of the button barcode model to give correct homolog pairing. All of these simulations, summarized in **Figure 3**, used buttons spaced at regular intervals in which the spacing between buttons was different on the two different chromosomes (details are provided in the figure legend) We model two sets of homologous chromosomes. We assess pairing fidelity in terms of the fraction of loci that are paired to loci on the correct homologous chromosome. Thus perfect fidelity would be reflected as 100% pairing to the correct homolog. For every chromosome, there is one correct homolog that it should pair with, but three incorrect chromosomes that it should not pair with (i.e. either of the two non-homologs also present, or else anywhere else on its own chromosome, i.e. pairing in cis). In the absence of any mechanism to promote pairing fidelity, one would expect to observe around 25% correct pairing just based on chance alone.

**Figure 3.**
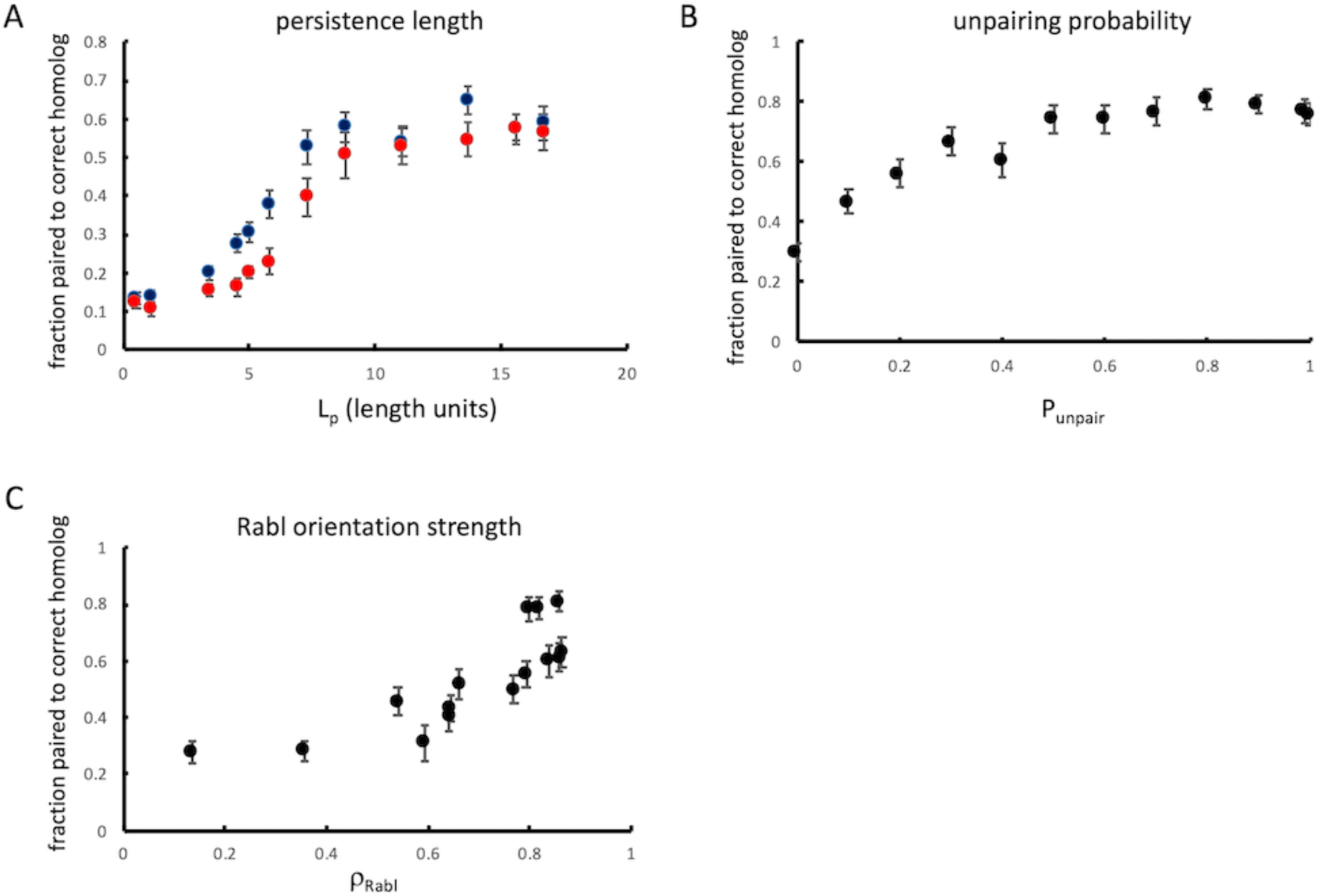
Influence of chromosome polymer physics and nuclear organization on homologous recognition. **A.** Effect of chromosome flexibility. Plot shows the results of simulations of pairing of two 5-button tracts with different spacing (3 vs 4 nodes between buttons), plotting the fidelity, defined as the fraction of nodes paired to their correct homolog, vs persistence length. The X axis is persistence length in simulation length units. Two sets of simulations were run, one in which the two tracts were centered on the same node in both chromosome pairs (blue), and the other in which the tracts were offset by 11 nodes between the two homolog pairs (red). All simulation results plotted are the average of 30 separate simulation runs. Error bars represent standard error of the mean. **B.** Effect of reversibility of pairing. Simulations were carried out using a uniform spacing between buttons (3 vs 4 nodes between buttons on the two chromosomes) with buttons spanning the whole arm, but with different values of p_unpair, the probability that two paired buttons will become unpaired at each timestep. All simulation results plotted are the average of 30 separate simulation runs. Error bars represent standard error of the mean. **C.** Effect of Rabl orientation. Plot shows the results of simulations as in panel B, but in which the confinement radius for centromere clustering was decreased to reduce the strength of the Rabl configuration. The X axis is the Rabl correlation coefficient, defined as the correlation between genomic position (distance from centromere) and vertical position (along the z axis of the nucleus). All simulation results plotted are the average of 30 separate simulation runs. Error bars represent standard error of the mean.

Our model for chromosome dynamics treats the chromosome as a worm-like chain, which can be characterized by the persistence length, which specifies the length scale over which the orientation of the polymer becomes uncorrelated. For a freely-jointed polymer, with no restriction on the bending angle at each node in the chain, the persistence length would be half the link length of the chain (Grosberg and Khoklov 1994) corresponding to 0.5 length units. In the button barcode model, the higher the persistence length, the greater the energetic penalty for associating buttons with different spacings on their respective chromosomes, and therefore the greater expected fidelity of pairing. As shown in **Figure 3A**, this is indeed the case - as the persistence length is increased (see Methods), the fidelity increases up to a plateau value. On the other hand, as persistence length decreases, the fidelity also decreases, down to a minimum when the persistence length is that of a freely jointed random coil. For such a freely jointed random chain, the fraction of loci paired to the correct homolog is actually less than the theoretical minimum of 25%. We interpret this to mean that for a highly flexible chain, a given locus can pair not only with its correct homolog plus two incorrect homologs, but also to other loci on the same chain, thus giving it more incorrect options, and that for a random coil, self-association might become highly favored due to the more compact shape of the folded coil compared to a more extended worm-like chain.

We also compared pairs of tracts that were located at the same distance from their respective centromeres, with pairs of tracts that were offset with one located near its centromere and the other located near its telomere (blue versus red markers in **Figure 3A**). Both arrangements showed the same general trend that increasing persistence length increased pairing fidelity, but higher fidelity was seen when the tracks were offset, which we interpret as an outcome of the Rabl configuration, which would tend to disfavor association of buttons located at different distances from their centromeres.

### Influence of pairing reversibility on pairing specificity

We next consider the influence of unpairing probability on the achievement of high fidelity pairing. Previous simulations of a specific button model (Child 2021) considered only irreversible pairing, which is appropriate for a model in which the on-rate for pairing is highly selective, to the extent that only correct (i.e. corresponding to identical loci on the correct homologs) buttons are allowed to pair in the first place. In our model, however, since button associations are completely non-specific, irreversible pairing would be expected to lock in incorrect associations, preventing them from ever being corrected. We therefore carried out simulations using a range of values for the parameter p_unpair, which describes the probability that two paired buttons might become unpaired during one iteration of the simulation. We have previously shown, in models with selective pairing sites, that high values for unpairing probability can still allow homologs to form stable associations (Marshall and Fung 2019). Here we investigate the effect of unpairing on the non-specific button barcode model. **Figure 3B** shows the fraction of loci paired to the correct homolog as a function of p_unpair. The worst performance is when p_unpair is zero, that is, when pairing is irreversible. This result is consistent with our intuition that reversible pairing would be needed to correct errors in association, and also matches our observations on simulations in which many incorrect associations can be seen at early time points, which are later corrected (see for example **Figure 2D** second time frame). In these simulations, maximum fidelity of pairing is achieved at a value of p_unpair of 0.8, but the actual maximum fidelity is a function of the specific barcodes chosen. Further increase in unpairing leads to a decrease in pairing, but it is interesting to note that even at a value of 1.0, meaning that every pair of loci will unpair at each iteration, homologous association still takes place. The reason for this effect is that when two buttons unpair, they remain near each other, and are thus biased to rapidly re-associate in the next time point. In any case, the main conclusion is that reversibility of association is not only tolerated, but is actually essential for selectivity in homolog recognition by non-specific buttons.

### Influence of large-scale nuclear architecture on pairing specificity

Chromosomes are not, in general, randomly arranged in nuclei (Comings 1980; Brickner 2017). One of the best -understood aspects of nuclear organization is the Rabl orientation, in which interphase chromosomes retain a vestige of their orientation from anaphase, such that the centromeres co-localize at one end of the nucleus, and the telomeres at the other, with the chromosome arms stretching in between. This configuration is seen in many different species and cell types (Comings 1980; Vourc’h 1993; Croft 1999; Cremer 2001; Carvalho 2001), but is perhaps most dramatically seen in *Drosophila* embryos (Marshall 1996; Dernburg 1996), where centromeres cluster at one end of the nucleus closest to the embryo surface and telomeres are at the other end of the nucleus.

One effect of the Rabl orientation is that corresponding loci on homologous chromosomes will tend to be nearer to each other than randomly chosen non-homologous loci, because they are the same genomic distance from their corresponding centromeres. Referring to the centromere-telomere axis as the vertical axis of the nucleus, these loci can be said to have similar vertical positions. This might give them a stronger tendency to associate with each other than with loci at other genomic locations, which would thus lie at different vertical positions.

In order to test the effect of the Rabl configuration on pairing fidelity, we performed a series of simulations using button tracts spanning the whole chromosome arm, with a spacing of 3 nodes between buttons on one chromosome and 4 on the other, in which we progressively reduced the degree of centromere clustering by increasing the diameter of the region in which the centromeres were confined. For each confinement region, we first ran the simulation without pairing and calculated the correlation coefficient between position on the chromosome and position along the z axis, which we take as a measure of the strength of the Rabl configuration. We then performed pairing simulations and plotted the average pairing fidelity versus the strength of the Rabl configuration. As shown in **Figure 3**C, the best pairing was obtained with the strongest Rabl orientation, and when the Rabl configuration was reduced to the point that chromosomal position and vertical position were uncorrelated, the pairing fidelity dropped to near the theoretical minimum value of 25%.

### Homology recognition using randomly generated button codes

Thus far, we have only considered the case in which different homologs have different, but uniform, spacing between their non-specific interaction “buttons”. We have shown that such a scheme can indeed lead to a majority of chromosome loci associating with the correct homolog compared to an incorrect homolog on which the buttons have a different spacing. But this scheme was arbitrarily chosen as a proof of concept, and we have no reason to believe it is the best possible way to achieve homolog discrimination. For one thing, by having regular spacing between all buttons on a chromosome, there is a potential for the pairing to get out of register, such that button n associates with button n+1 on the homolog. Such out of register association with the correct homolog would not incur an energetic penalty in that neither homolog would be required to stretch or bend to achieve alignment. A second limitation of regular spacing is that different regular spacings on different chromosomes requires different densities of pairing sites on different chromosomes, which may or may not be biologically desirable. More generally, if we want to use our theoretical work as a source of hypotheses about the possible distribution of pairing sites in actual chromosomes, it is important to have an idea of what pairing site distribution is optimal, under the assumption that selection pressure may have driven a similar distribution in real chromosomes.

In order to see how the pattern of buttons along a chromosome might influence the fidelity of homolog recognition, we generated a series of random button distributions and simulated pairing in each case. As shown in **Figure 4**, when we generate random codes at three different densities of buttons, we observed a range of pairing fidelity. The majority of random codes were able to give pairing fidelity greater than 60%, which is comparable to the level of correct somatic homolog pairing in many actual cases (see Discussion). In one case, the fidelity was extremely high (>99%). The fact that almost perfect homolog pairing could be obtained just by sampling a small number of random codes suggests that a button barcode could be easy to evolve.

**Figure 4.**
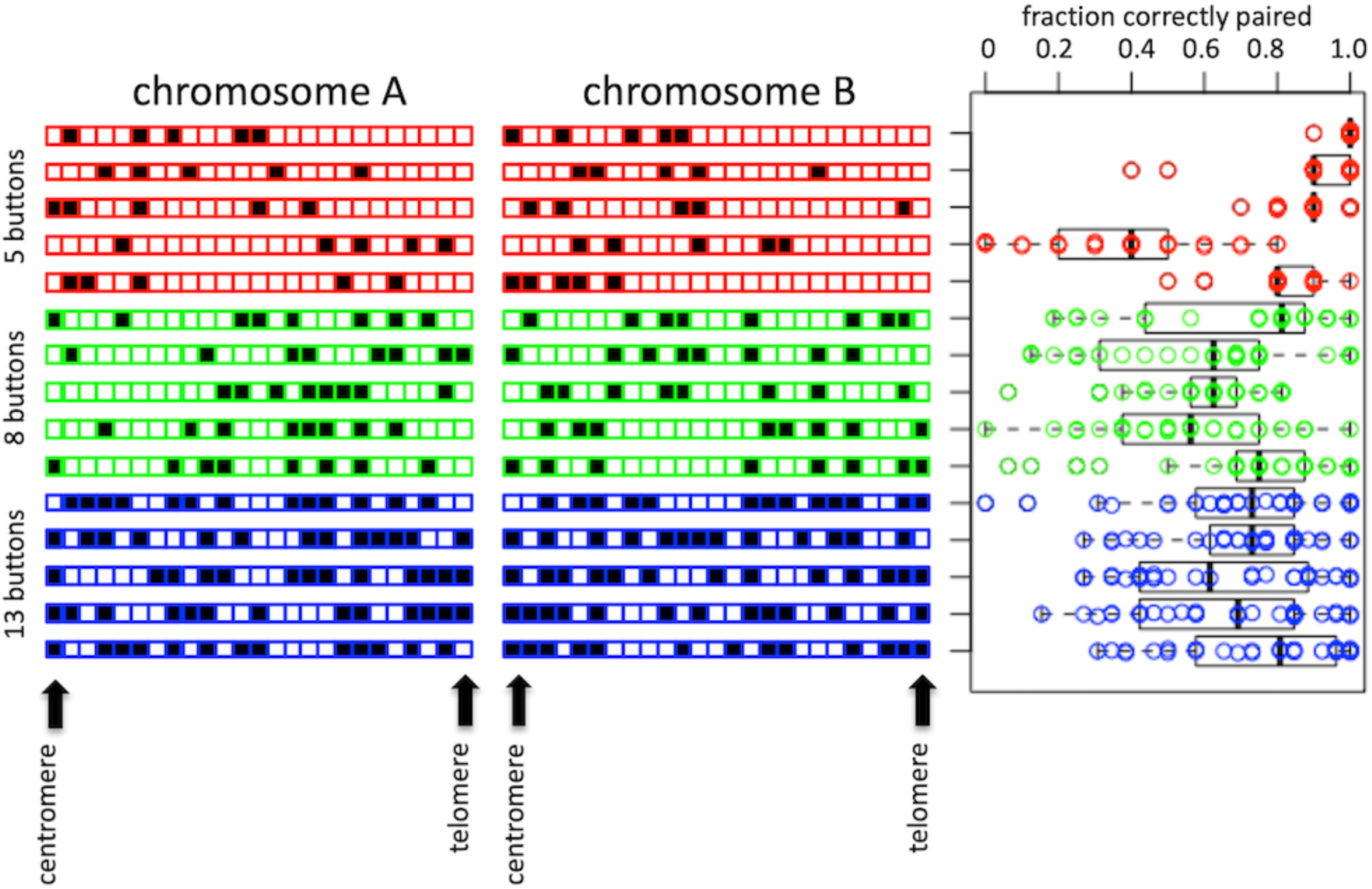
Randomly generated coding can achieve selectivity. Bar diagrams show the location of pairing buttons on the two chromosomes, with each square corresponding to a node on one chromosome arm, and a block box corresponding to a node with a pairing button. Red, green, and blue illustrate randomly generated arrangements of 5, 8, and 13 buttons, respectively. The Beeswarm plot on the right shows the pairing outcomes (fraction of loci paired with the correct homolog) for 30 simulations of each random code pair. In each case, the theoretical minimum pairing fidelity expected, if all buttons were pairing non-selectively and independently of each other, is 0.25.

Looking at the five-button random codes, the code pair that gave the best performance was one in which the buttons on both chromosomes were clustered near their corresponding centromeres. We hypothesized that part of the reason these codes worked so well might be that their proximity to the centromere makes them maximally subject to the Rabl constraint, since in our model this constraint was implemented solely by clustering centromeres together. To test this idea, we repeated the simulation of the same pattern of buttons, shifting the pattern progressively away from the centromere. As shown in **Figure 5A**, the fidelity of pairing decreased continuously as the tracts were moved away from the centromeres, consistent with the idea that the button barcode segments work best when located near clustered centromeres.

**Figure 5.**
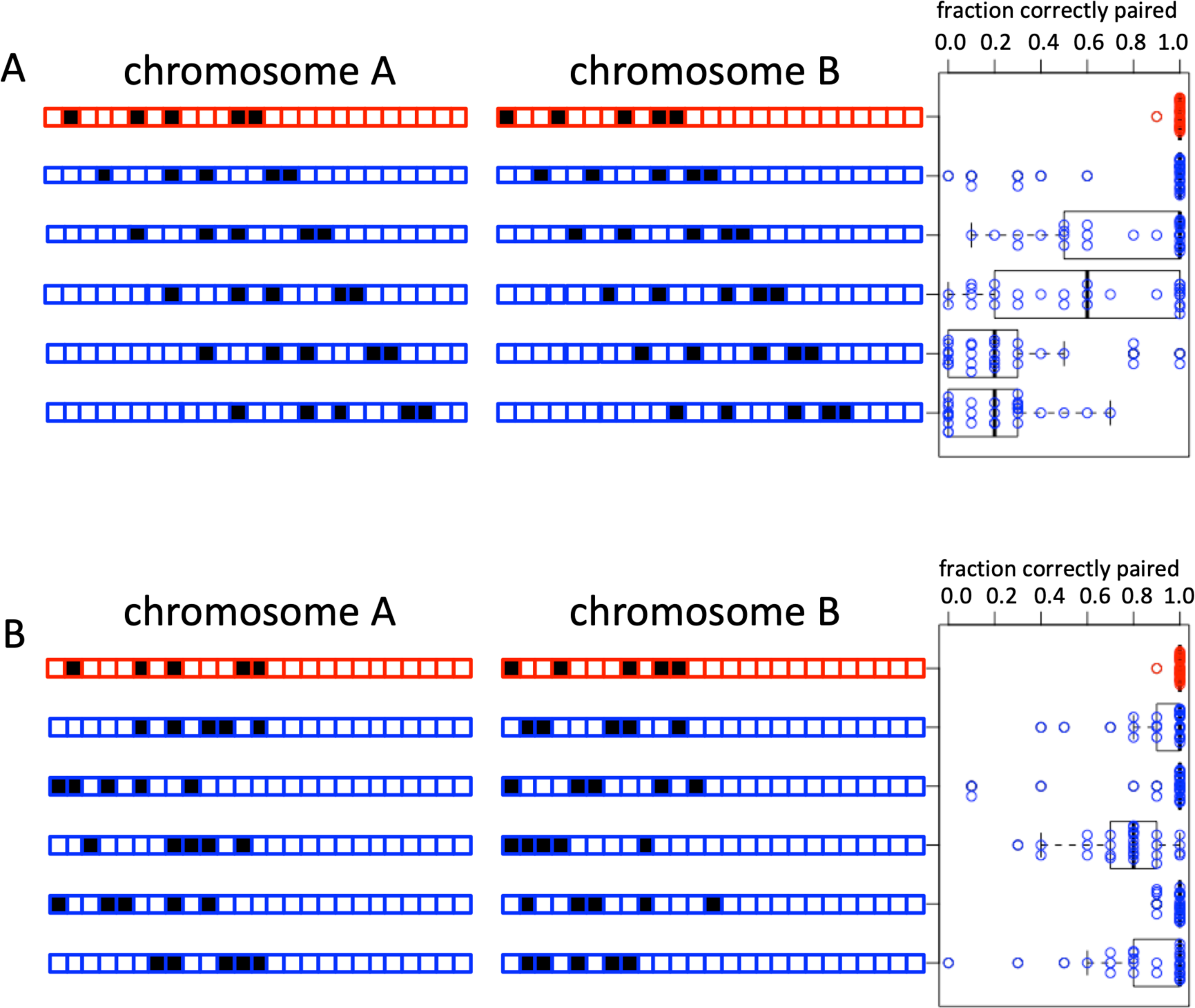
Variations on the 5 button randomly generated code. **A**. The best-performing randomly generated code from Figure 4 was shifted progressively away from its centromere in intervals of two nodes. The red bar illustrates the original location of the buttons. For each shifted version, the bar plot displays the location of the buttons. The beeswarm plot shows the corresponding pairing outcomes from 30 simulations for each code pair, illustrating degradation of performance as the button segments are shifted away from the centromeres. **B.** Testing other randomly generated five button codes that are constrained to span the same range of nodes as the optimal five button code from Figure 4 (which is shown in red). Four of the five random code pairs give essentially as good pairing outcomes as the original code pair.

Based on these results, we generated a further set of random five-button barcodes, this time constraining them to all lie within a 13 node stretch of the simulated chromosome at the end near the centromere. As shown in **Figure 5B**, two of the five additional random 5 button codes were also highly effective for chromosome pairing, comparable to the original high-performing random 5 button code from **Figure 4**.

We conclude that random arrangements of nonselective buttons positioned with non-uniform spacing can in fact lead to extremely efficient homology recognition. However, among the randomly generated non-uniform button patterns, the ones that work the best seem to be those that are restricted to sub-regions of the chromosome arm.

In an effort to obtain effective codes that span the whole arm, we next turned to actual barcodes encountered in everyday life.

### Homology recognition using an industrial barcode

Barcodes are familiar in our everyday lives, printed on virtually all commercial products. Barcodes are patterns of black vertical stripes separated by white vertical spaces (Palmer 1995). Using just these two colors, it is possible to discriminate a large number of different symbols, based on the pattern of widths of the stripes and spaces (Pavlidis 1990). Most bar codes, such as UPC or Code 39, encode information in the widths of both the white and black bars. This is fundamentally different from the chromosome pairing code we propose here, in that in the chromosome pairing case, only the spacing between the pairing sites encodes information, and all the pairing sites themselves are treated as equivalent. This would be analogous to a barcode in which all the information is encoded by the widths of the bars, but all spaces have equal length. In fact, such barcodes do exist, the most common example being code 2 of 5 (Palmer 1995).

A second feature of real barcodes is that in general the width of the stripes is constrained to take on just one of two values, such that there are wide stripes and narrow stripes and no other options. For most bar code symbologies, the wide stripes are three times wider than the narrow stripes, with the widths selected so as to maximize the difference between wide and narrow stripes, while subject to constraints concerning the minimum size of the narrow stripe and the total number of stripes in the coded character (Palmer 1995). In a few rare examples, such as the Codabar barcode, the bars are not multiples of a unit width and multiple different bar widths end up appearing within a code (Allais 1984), however this variation in bar widths ends up not increasing the information capacity, and such codes can be replaced with variants using just two bar widths (Harmon and Adams 1989.

A third feature of real barcodes is that the order in which the bars occur is critical - if the bars are read in a different order, the symbol will be decoded differently. This is generally not an issue in real barcodes that are printed onto a rigid surface, but for a chromosome bar code, there is certainly the potential for bars (which we now interpret as the spacing between successive pairing sites on a chromosome) to be read out of order, depending on how the chromosome polymers are folded. Our data above showed that reliable decoding (i.e. choosing the correct pairing partner) requires that chromosomes are not just random coils, but maintain some linear structure due to the physics of a worm-like chain. The more rigid the chain, the more the arrangement of spacings between pairing sites will approximate a real barcode printed on a solid surface.

Finally, during the decoding of a real barcode, the symbol is scanned from one end all the way to the other. This allows the bars to be read in the correct order, which essentially means that as each bar comes up, it can be compared computationally to an internally stored reference pattern. In the case of the chromosome button barcode, decoding the bars in the correct order is enforced by the Rabl orientation.

There is thus a very concrete analogy between real bar codes and chromosome pairing barcodes, suggesting it would be possible to encode an actual barcode on a chromosome (**Figure 6A**). To do this, we start with the 2 of 5 code (Allais 1984) in which (a) all information is encoded by the widths of the black bars, (b) there are just two possible widths for the black bars, with the wide bars being three times as wide as the narrow bars, (c) all white bars are the same width corresponding to the narrow black bars, and (d) every character is encoded by five bars, of which two are wide. This code was designed to represent the digits 0-9 (**Figure 6B**). To generate a chromosome pairing code based on 2 of 5 code, we treat the pairing buttons on a chromosome as the white bars in code 2 of 5, and the gaps between pairing buttons as the black bars. To achieve the 3:1 ratio of wide to narrow stripes in code 2 of 5, while spanning most of one chromosome arm, we space buttons 6 nodes apart for a wide bar and 2 nodes apart for a narrow bar. **Figure 6C** shows the chromosome implementation of two different digits, 0 and 1, in 2 of 5 code. As seen in **Figure 6C**, chromosomes printed with 2 of 5 code can be discriminated with a reliability of 0.91.

**Figure 6.**
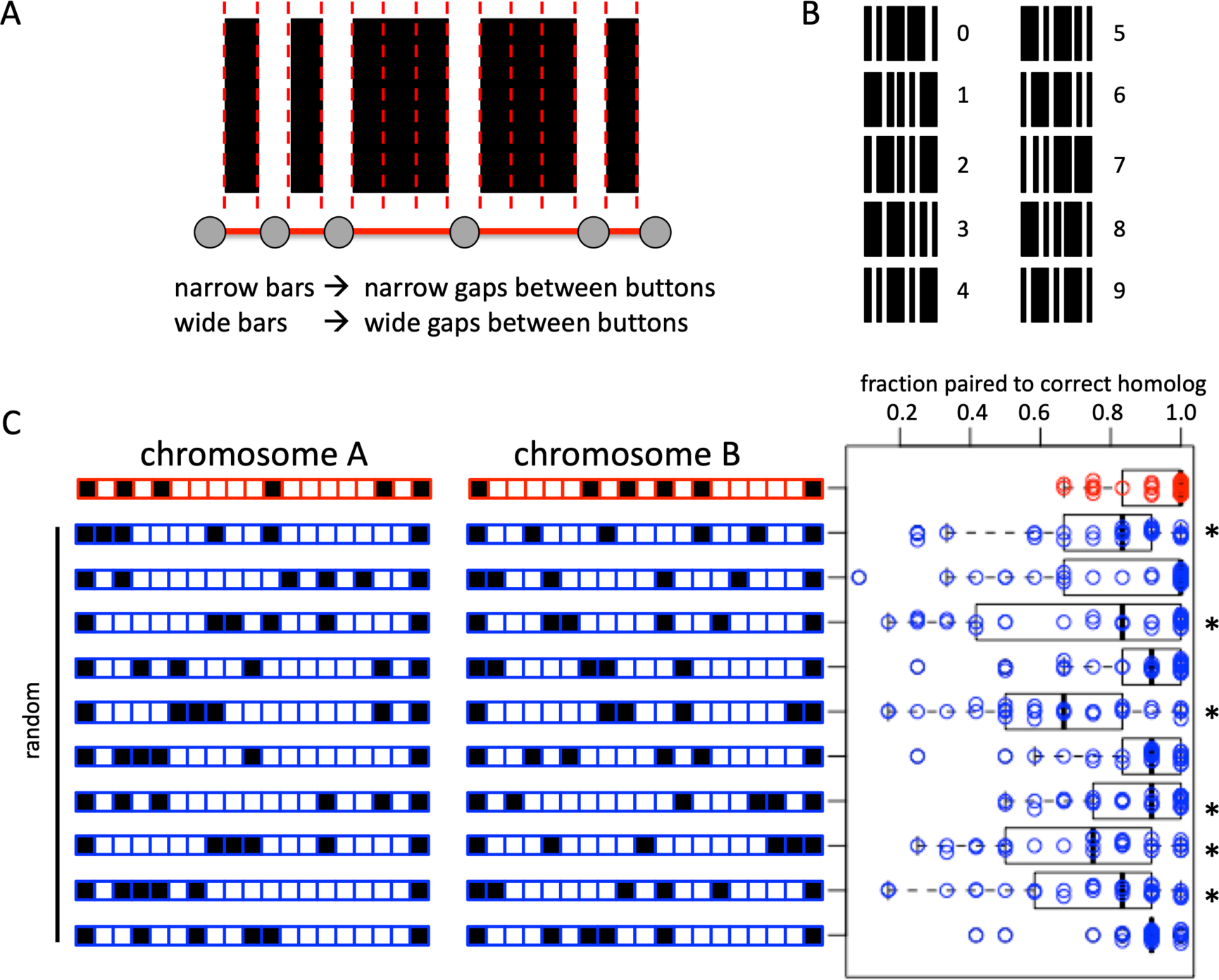
An industrial barcode can achieve selectivity for homolog pairing. **A.** Analogy between an industrial barcode and the pattern of chromosome button spacing. Buttons on the chromosome correspond to the white bars in the industrial bar code, which are all identical. Gaps between buttons correspond to the width of the black bars in the industrial barcode, which take on two values, narrow and wide. **B.** The “code 2 of 5” barcode symbology (Harmon and Adams, 1989). In this barcode scheme, information is only encoded in the width of the black bars, while the spacing between the bars carries no information. Every character consists of five bars, two wide and three narrow. **C.** Button barcode derived from 2 of 5 by setting each narrow bar equal to a gap of two nodes between buttons, and each wide bar equal to a gap of 6 nodes between buttons. Beeswarm plots on the right show the outcome of a 2 of 5 code using symbols for 0 and 1, compared to random simulations with the same number of pairing sites distributed with the same endpoints. Asterisks denote random codes which gave significantly less correct pairing compared to the 2 of 5 barcode simulations, at a significance of 0.05 or better based on a one-tailed Mann Whitney test.

For comparison, we also simulated random codes with the same number of buttons as the 2 of 5 code and spanning the same range of nodes, with buttons fixed at the same endpoints. While several of the random codes performed almost as well as 2 of 5 in terms of their average fidelity, the 2 of 5 barcode simulation showed a clear difference in terms of the left tail of the distribution - in comparison to the random codes which sometimes gave very poor results, 2 of 5 never had less than 60% match to the correct homolog. This suggest that the industrial barcode performs better than comparable random patterns in terms of minimizing worse case results.

## Discussion

### Comparison with pairing levels reported in other studies

In our simplified model, we find that non-specific associations can result in specific pairing frequencies exceeding 90%. However, our model does not reliably achieve 100% pairing for any parameter values we have tried. Our model is not intended to represent any specific actual species or cell type, but rather to be an abstract model to test the general concept of button barcodes. Nevertheless, the fact that the model cannot achieve 100% pairing raises the question of how this compares with what is seen in actual cells?

Chromosome-wide somatic pairing is most notable in *Drosophila*. A FISH survey of 11 loci on the left arm of chromosome 2 in cycle 14 *Drosophila* embryos found a range of pairing frequencies from 7% to 85%, with most loci in the range of 20-30% paired (Fung 1998). The same study found that later in embryonic development, by 6 hours AED, pairing frequencies increased to 20-98%, eventually reaching 80-100% by day 5 of development. Other FISH analyses in *Drosophila* embryos gave pairing frequencies of 60-90% for cycle 14 (Hiraoka 1993) and 70% in post-gastrulation embryos(Gemkow 1998). In the *Drosophila* eye, Viets (2019) found that pairing loci associated at frequencies in the range of 88%-94%. Screening studies of specific loci under a range of perturbations gave control levels of pairing in the range 47-91% for *Drosophila* embryos (Bateman and Wu 2008) and 40%-80% for *Drosophila* Kc167 cells (Puerto 2023).

In human cells that show pairing of only certain chromosome regions, rather than whole chromosomes like in *Drosophila*, pairing frequencies have been reported to be in a similar range (e.g. Arnoldus 1989).

It is thus clear that while our model does not achieve 100% pairing efficiency, neither do actual chromosomes. We conclude that the level of pairing achievable even with a simple model based on non-specific associations can, at least in principle, produce the necessary level of correct pairing.

We note that our analysis, as well as all existing data of which we are aware, has focused on the faction of loci that pair with their correct homolog, as the primary figure of merit. Since we do not know the physiological function (if any) of somatic homolog pairing, it seems reasonable to assume that the system has evolved to maximize pairing with the correct homologous chromosome. It is also possible that chromosomes have been selected to avoid pairing with incorrect chromosomes. From an experimental point of view, it is unclear how to detect incorrect pairing. Correct pairing can be assessed by probes that label specific loci, because for correct pairing, the two loci involved in the interaction are known, so that probes can be designed for either fixed or living cells. But an incorrect pairing could in principle take place anywhere in the genome, and given our proposed non-specific button interaction, these incorrect pairings would involve different loci in every cell. Thus, while methods such as Hi-C can reveal all near spatial contacts between chromosomes, and can be modified to detect associations between homologs (Erceg 2019), to our knowledge there would still be no way to distinguish an actual incorrect pairing from a fortuitous proximity of two loci. We thus propose that there is a need for methods to define and detect incorrect pairing events. The challenge is to have a way to distinguish actual pairing from fortuitous random contact. This could be done in live cells by tracking correlated motion of two loci, it is not obvious how to perform such an analysis on a genome-wide scale.

### Specific versus nonspecific pairing buttons

We have posed our nonspecific button barcode model as being fundamentally different from a specific button model (**Figure 7A,B**), but this distinction may not actually be so clear. We have shown in our simulations that while barcodes spanning an entire arm can achieve specificity, we also find that even short barcodes involving just a few nonspecific buttons can also pair selectively (**Figure 5B**). This suggests a hybrid model (**Figure 7C**), in which short barcode segments of nonspecific buttons can selectively associate with each other based on the pattern of spacing between their button elements, but then these barcode segments would serve as selective buttons for overall chromosome pairing. With this picture in mind, we consider the existing evidence about the nature and specificity of pairing buttons in *Drosophila*.

**Figure 7.**
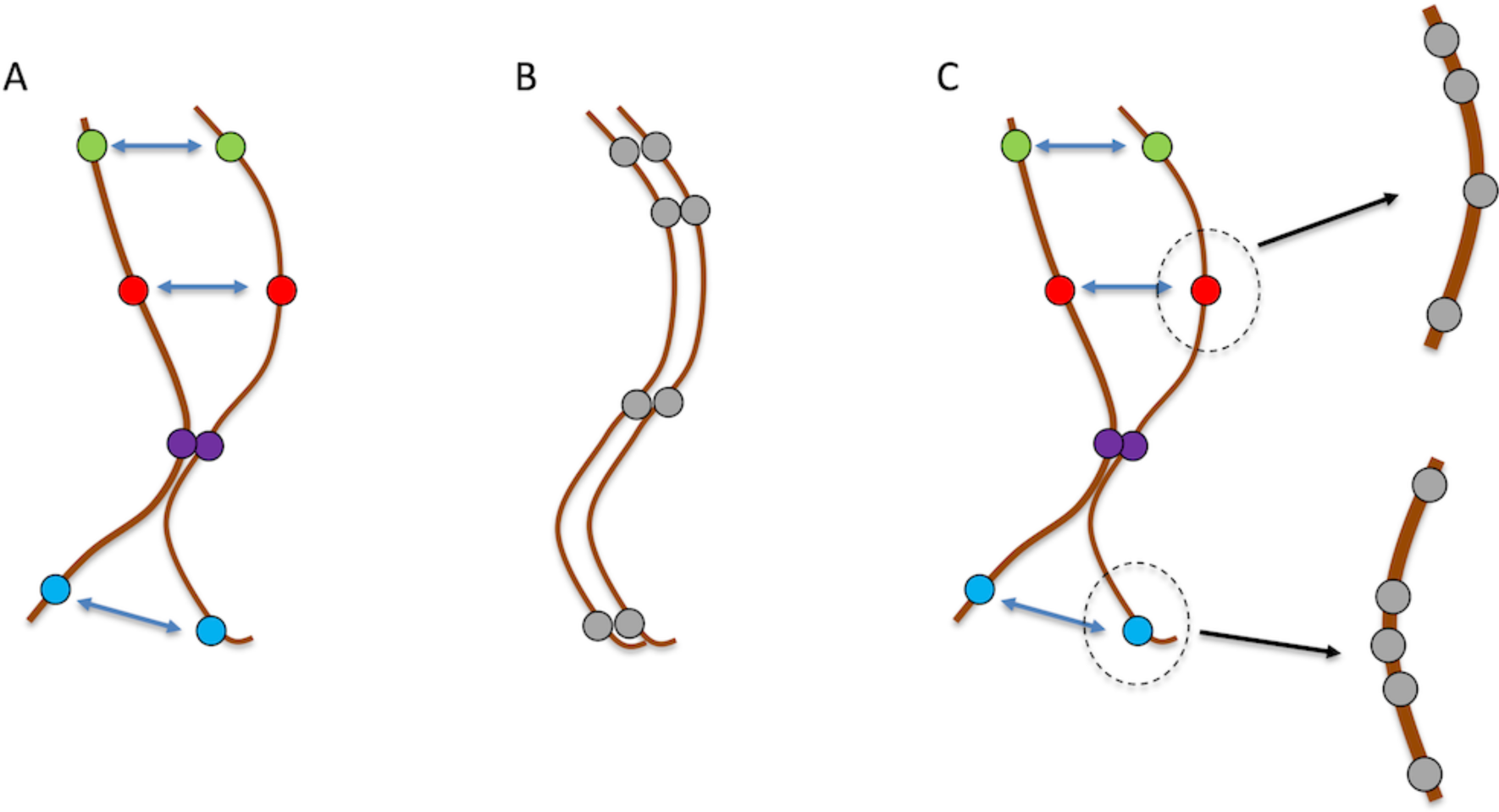
Button models for homolog recognition. **A.** Specific Button model, in which each pairing button can only associate with an identical type found on the homolog, as indicated by the different colors. These types of models have been previously discussed and in some cases analyzed by computational modeling (see for example Fung 1998; Viets 2019; Child 2021). **B.** Non-specific Button Barcode model, in which all pairing buttons can associate with each other with the same intrinsic affinity, but the non-uniform spacing between pairing buttons on different chromosomes creates a barcode effect to give specificity. **C.** Barcode Patch model, in which short segments containing non-specific association domains arranged in distinct bar-code patterns creates larger scale specific buttons, without at any point requiring homolog specific molecular interactions.

The model presented here predicts that large-scale chromosome rearrangements would strongly disrupt homolog recognition. For example, a translocation would delete part of one bar code and add it to a different bar code on another chromosome. A deletion would alter the number of buttons or the spacing between them while a duplication would add buttons or increase the spacing. Indeed, a model based on a barcode of nonspecific buttons should be much more sensitive to the effects of chromosome rearrangements than a model based on specific buttons, which would retain their specificity if moved to another chromosome. Thus, to distinguish a button barcode model from a specific button model, one key class of experiments would be analysis of the effect of chromosome rearrangements on pairing.

Much of the genetic information about translocation effects on pairing in *Drosophila* come from studies of transvection, in which the enhancer of one gene can regulate expression of its homolog, presumably involving somatic pairing. There is an extensive literature showing that chromosome rearrangements can disrupt transvection, presumably via an effect on pairing, and that this effect tends to be largest when the translocation breakpoint is in between the gene and its centromere (Duncan 2002). A disruption of homolog pairing at one specific locus, the histone locus on chromosome arm 2L, was directly observed by FISH for the ltx13 translocation in *Drosophila*, which moves most of the left arm of chromosome 2 onto the end of the right arm of chromosome 3. Somatic pairing of the histone locus on 2L is almost completely lost when this translocation is heterozygous, but is mostly restored if the translocation is homozygous (Hiraoka 1993). There is thus a clear set of cases in which chromosome rearrangements do indeed disrupt pairing.

This seems to be consistent with the button barcode model but an alternative interpretation is that the translocations that affect pairing at a given gene could work not by affecting pairing per se, by separating that gene from a nearby specific pairing button. In this interpretation, the buttons are still paring normally, but because the gene being tested by transvection or FISH isn’t actually inside the button, it loses its own pairing when separated from the button by a translocation. An alternative strategy to looking for a loss of pairing caused by a translocation breakpoint, is to insert an ectopic copy of a genomic region elsewhere in the genome and ask if this is able to confer pairing to the original site. This approach was taken by Viets (2019) who identified a number of sites with this property. These regions, capable of ectopic pairing when relocated in the genome, were on the order of 100 kb in size. The whole region was required - when smaller sub-regions were tested, they did not have the ability to pair. But not every region could confer pairing when relocated, consistent with a specific button model, in which the buttons are not individual DNA elements such as enhancers or insulators, but larger regions on the size scale of TADs.

Genetic analysis of transvection at different genes has shown that pairing is only affected when breakpoints are within some distance of the gene in question, but the distance depends on the locus. For some genes like Ubx, any translocation breakpoint over a very large region can disrupt transvection (Lewis 1954), while for other genes such as yellow and white, only translocations within a few hundred kb of the gene have such an effect (Smolik-Utlaut 1987; Ou 2009). The size of the regions for yellow and white is comparable to the size of the buttons identified by Viets. Taken together, these data suggest that for *Drosophila*, a model based on specific buttons, rather than a nonspecific button barcode, may best explain the data on translocation effects.

Nevertheless, the question remains of how specificity is achieved within the hundreds of kb pairing regions. Based on our model, we propose that it might be the spacing between smaller elements within such a domain that determines specificity, in a bar-code scheme like that proposed in the present study. In our models above, we showed that a short series of buttons could associate chromosomes together at one point, and we suggest that the buttons described by Viets may correspond not to single buttons in our model, but to short patches of buttons arranged in specific barcode patterns. We note that the spacing between the non-specific interaction elements within a TAD would be substantially smaller than the distances between buttons in our model. In fact, for a bar-code spanning a much smaller length like a single TAD rather than a whole chromosome, the distances between elements would be less than the persistence length of the chromatin, hence the polymer would behave even more like an elastic rod, making the bar-code mechanism even more effective. This model is similar to the insulator code model proposed by Viets, in which buttons are recognized by having different combinations of insulator elements, except that in the barcode patch model, the individual pairing determinants within the button (or TAD) would no longer need to be different from one another or to have any specificity, and differences in their spatial arrangement would be the source of specific association.

One can therefore imagine two types of “buttons” existing at two different scales, in which at the level of the whole chromosome, pairing is apparently mediated by specific buttons as per the Viets (2019) model, but in which these large scale buttons actually represent barcode patches comprising non-specific interaction buttons (**Figure 7C**). Thus, the ultimate source of specificity would still be a bar code of non-specific interaction buttons, but these would exist at a smaller scale relative to the whole chromosome. This type of model could therefore explain the ability of short translocations to confer pairing as per Viets (2019) but would not explain the ability of large-scale translocations to disrupt pairing (e.g. Hiraoka 1993). In the latter case, the effect could be dominated not by loss of bar-code recognition, but by the fact that the large translocations result in two pairing patches being localized in distinct regions of the nucleus where they never get the chance to interact (Marshall 1996; Erceg 2019).

### Implications of chromatin physics and cell biological constraints for homology recognition

The ability to achieve specific homolog recognition in the non-specific button barcode model depends on the mechanical properties of the chromosomes and their arrangement in the nucleus. **Figure 3** indicates that specificity is strongly influenced by the persistence length of the chromatin, by the parallel arrangement of chromosomes caused by the clustering of centromeres at one end and telomeres at the other (Rabl orientation), and by the reversibility of associations between homologs. Are the physical properties of actual chromosomes in an appropriate regime for this type of model to work? The importance of pairing reversibility has been previously discussed in the context of meiotic chromosome pairing (Kleckner and Weiner 1993) and has been directly observed during meiosis in living yeast cells (Newman 2022). It is therefore plausible that pairing for somatic chromosomes would be similarly dynamic.

Here, we consider persistence length. In our model, we simulated persistence lengths in the range from 0.5 to 16.7 length units, corresponding to 0.1 - 3.3 microns. We found that correct associations occur frequently when the persistence length exceeds roughly 5 length units, which corresponds to 1 micron (**Figure 3A**). This is much longer than the persistence length of 50-80 nm for DNA reconstituted with nucleosomes (Garai 2015), but is only several fold higher than the persistence length of 220 nm reported for yeast interphase chromosomes (Bystricky 2004). Our maximum persistence length is on the same size scale as the nucleus itself, but is still about 50 fold smaller than measurements of persistence length of mitotic chromosomes in *Drosophil*a (Marshall 2001). The persistence length are therefore consistent with the extension of chromosomes from one end of the nucleus to the other in the Rabl orientation, as well as with the fact that in the rapid cell cycles of *Drosophila* early embryos, chromosomes may not fully de-compact.

The key question with respect to persistence length in the model is whether, on the length scale of spacing between pairing sites, the chromosome is better treated as a random walk polymer or an elastic rod. If the persistence length is short compared to the pairing site spacing, the chromosome would behave like a random polymer and the average distance between sites on a chain in 3D space would be proportional to the square root of the length along the chain. If the persistence length is long compared to the pairing site spacing, the chromosome would behave like an elastic rod and the average distance between sites on the chromosome in 3D space would be proportional to the length along the chain. To decide which regime applies in the case of *Drosophila* embryos during the time of somatic homolog pairing, we refer to a prior analysis of nuclear position of loci spanning the left arm of Chromosome 2 in cycle 13 *Drosophila* which showed that position along the chromosome was highly correlated with position along the nuclear axis (Marshall 1996), confirming the presence of a strong Rabl orientation in *Drosophila* embryonic nuclei. Replotting that data by averaging positions for loci in a given segment of the arm (each arm of a *Drosophila* chromosome has 20 cytologically defined segments), and then plotting vertical position versus genome position for eight segments of the left arm of chromosome 2, we find that vertical position is well fit by a linear function (**Supplemental Figure 1**), suggesting that the interphase chromosome is behaving more like an elastic rod than a random chain, and suggesting that the persistence length should be at least on the order of the size of the nucleus, which fits with the 1-3 micron range in which we see effective homolog recognition in our model.

The Rabl orientation was found to promote pairing selectivity in our model (**Figure 3C**). This parallel alignment of chromosomes tends to place homologous loci in similar regions of the nucleus, but also ensures that the buttons along one chromosome will tend to align with buttons located in the same linear order on other chromosomes. From a barcode perspective, this is analogous to reading the bars of a barcode in order. In the absence of a Rabl orientation, chromosomes could intersect at various angles and it would be possible, for example, for one set of buttons to align with another set in reverse order. The Rabl orientation is thought to largely arise from the face that centromeres after mitosis remain clustered together at one end of the nucleus. In our model, we found that short barcodes gave better pairing fidelity when they were located near the centromeres, suggesting that the influence of the Rabl orientation is strongest near the point of clustering. This is consistent with studies in yeast showing that the influence of telomere tethering to the NE on chromatin position is strongest for loci located near the telomeres, and drops off as one looks at loci located farther away on the chromosome (Avşaroğlu 2014).

### Potential implications for meiotic homolog pairing

Our study reported here focuses entirely on somatic homolog pairing, where recombination does not occur and no mechanism is known for DNA strand invasion or annealing. Given that DNA contains sequence information, and given the involvement of recombination and strand invasion as a core part of meiosis, it seems obvious that sequence comparison would be the basis for homologous recognition in meiosis, thus suggesting a specific button model in which buttons correspond to double strand breaks and their corresponding homologous DNA sequences. Genetic studies have found that mutations affecting recombination impair homologous association during meiosis in yeast and mice (Keeney 1998; Romanienko 2000), supporting the idea that a DNA-base homology level search is at work.

Sometimes, however, recombination-independent homologous association is seen even in meiotic cells. For example, in male meiosis in *Drosophila*, double strand breaks and recombination do not take place, and yet homologous chromosomes still associate (McKee 2012; Rubin 2022).

In *C. elegans* (reviewed in Rog and Dernburg, 2013), homolog pairing does not require recombination (Dernburg 1998), and instead, pairing is dictated by chromosome segments known as pairing centers (PCs). When PCs are deleted, pairing is eliminated, and when they are translocated to another chromosome, pairing becomes dictated by the new PC (McKim 1988; Zetka and rose 1992; MacQueen 2005). These PCs are needed not only for pairing but also for interaction of the chromosomes with the cytoskeleton through the nuclear envelope, thought to be important for establishing proper nuclear organization and driving active chromosome motion (Rog and Dernburg 2013). Each PC contains a large number of short sequence elements that bind a set of zinc finger proteins known as ZIMs. Each PC recruits a single type of ZIM (HIM-8 for the X chromosome, ZIM-3 for chromosome 1, etc.), but in some cases, the same ZIM is recruited to more than one PC (for example ZIM-3 is recruited to both chromosome I and chromosome IV, while ZIM-1 is recruited to the PCs for chromosomes II and III). Artificial arrays of these zinc finger binding sequences can replace the normal requirement for a PC and can even cause ectopic pairing to non-homologous chromosomes when translocated into new contexts (Phillips 2009). These PCs seem like the most canonical possible example of a specific button code, in that each PC only associates with the homologous PC. The obvious model is that ZIMs recognize the sequences, and then interactions between the ZIMs drives pairing. But this cannot be the whole story because of the fact that more than one PC shares the same ZIM. Within each PC, the zinc finger binding sequences occur in clusters that are separated by long stretches of intervening DNA sequence (Phillips 2009). We propose that the differential distribution of zinc finger binding sequences along each PC provides specificity, even when the same ZIMs are used, using essentially the same barcode patch mechanism (**Figure 7C**) that we have described here.

In light of our results that the highest selectivity for a short barcode is seen when the code segment is near the centromere cluster, we speculate that part of the reason that nematode PCs need to be associated with the NE and cytoskeleton is to achieve a similar arrangement of chromosomes with all the barcode patches near the site of clustering.

There is also evidence that even in organisms such as mice and budding yeast, that rely on recombination for full pairing and synapsis, homologous chromosomes are already associated with each other prior to the onset of DSB-mediated pairing (Weiner 1994; Burgess 1999; Boateng 2013; Grey and de Massey, 2021; Solé 2022). It has been specifically shown that DSB formation by the SPO11 enzyme was not required for this pairing to occur (Boateng 2013). This pre-meiotic chromosome pairing might very well entail similar mechanisms as somatic homolog pairing, and we therefore speculate that non-specific button barcodes might potentially play a role. One potential candidate that has been proposed for such a role in meiotic chromosomes is cohesin (Ishiguro et al. 2014), but one could invoke a wide range of possible interactions mediated by proteins or other molecules associated with the chromosomes.

Even in cases where sequence level comparison mediated by meiotic DSBs is the sole source of specificity for pairing, the results presented here suggest that the spatial arrangement and polymer physics of chromatin could contribute to discrimination of correct homologous sequences from homeologous sequences present on other chromosomes. For one thing, in many cell types the telomeres cluster to form a bouquet, which has an effect similar to the Rabl orientation in mitotic cells, of aligning the chromosomes in a parallel fashion such that loci at similar distances from their corresponding telomeres will tend to occupy similar regions of the nucleus. For any given locus undergoing search via the recombination machinery, it will be less likely to make a false interaction with homeologous regions, even if they have high sequence similarity, if those regions are present at a different distance from the telomere. This is particularly likely in species like *S. pombe* or *Tetrahymena*, where the chromosomes are drawn out into long parallel “horse tail” configurations (Chikashige 1994; Loidl 2004). This stretching involves clustering of both telomeres and centromeres (Loidl 2012; Tian 2020) and is thus highly analogous to the Rabl configuration seen during somatic homolog pairing in *Drosophila*.

Chromosome elasticity also could play a role in conventional DSB-mediated meiotic pairing. The same physical effects in our model that favor pairing of non-specific buttons having similar spacings along their respective chromosomes, would favor a given pair of DSBs on one chromosome associating with homologous regions spaced similarly on the other chromosome. Even if each DSB was potentially capable of base pairing with several different homeologous sequence stretches elsewhere in the genome, these alternative regions will not in general be spaced the correct distance on the non-homolog, and thus false associations will be disfavored due to chromosome elasticity. It has been shown that mutation in cohesin leads to increased ectopic pairing in yeast (Lui 2013). We speculate that this decreased fidelity of homolog recognition might result from a reduction in the chromosome stiffness, such that it becomes more like a random walk, and therefore less able to benefit from the proposed mechanical barcode effect.

The idea that even when specific interactions are at work, additional discrimination is achieved by the physical “fit” of two interacting molecules, has a strong precedent in DNA replication fidelity, where even if an incorrect nucleotide can base-pair with the template strand via non-canonical base pairing, the geometry of the nucleotide in the active site is such that additional energy is required to deform the polymerase to accommodate the nucleotide, which creates a kinetic barrier to incorporation (Johnson 1993)

## Acknowledgments

We thank Eric Navarro and Sy Redding for discussions about how polymer physics may affect chromosome pairing dynamics. The authors acknowledge funding from NIH grants R35 GM130327 to WFM and R01 GM137126 to JCF, as well as support from the Center for Cellular Construction funded by NSF grant DBI-1548297.

**Note:** This is a preprint and is not peer reviewed

---

**Supplemental Figure S1.**
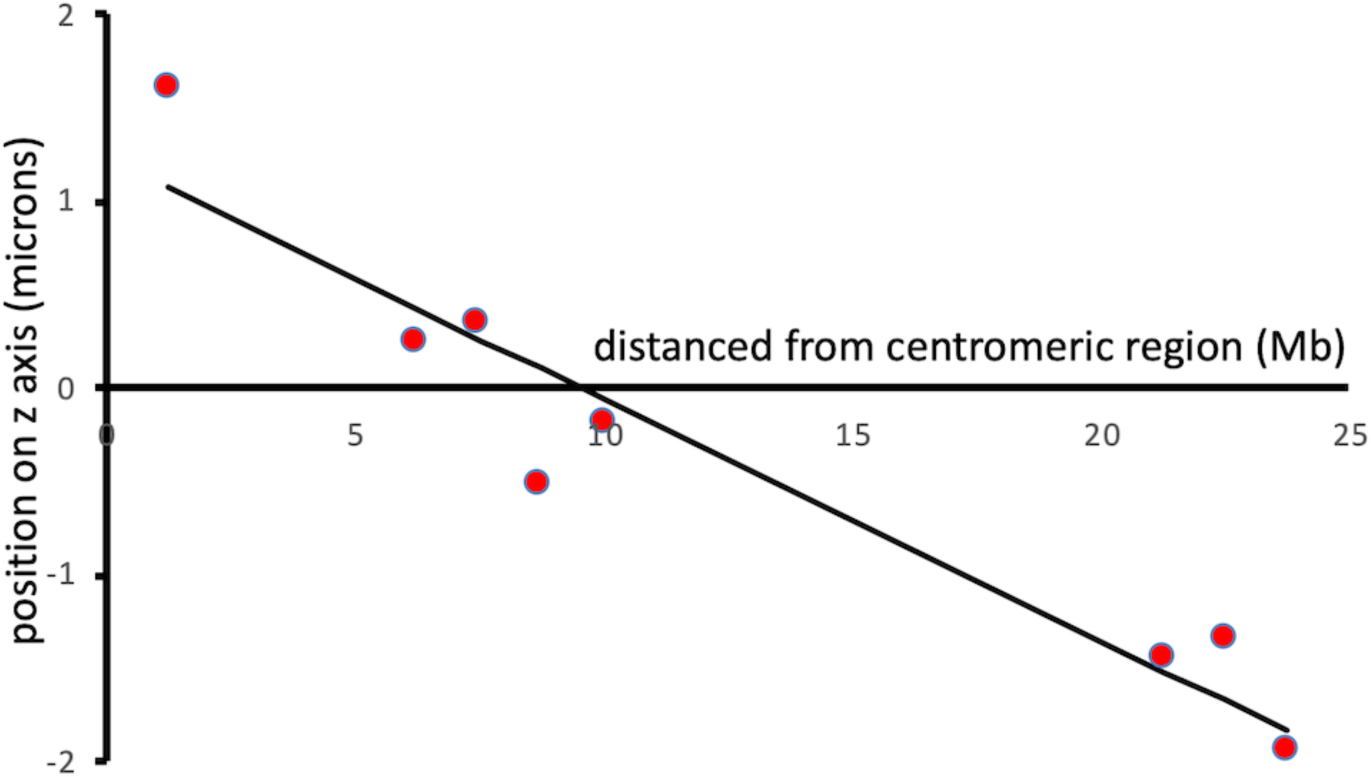
Fitting z axis position to genomic position in *Drosophila* embryos supports an extended worm-like chain rather than a random coil at the length scale of the whole chromosome. Previously published three dimensional FISH data from cycle 13 *Drosophila* embryos (Marshall 1996) were binned based on segment along chromosome arm 2L, and the average vertical distance (z) plotted as a function of the genomic distance of the segment from the centromeric region, in Mb. Best fit line is included with slope −1.3 microns per Mb. These data show that the chromosome arm spans a distance along the z axis of a little over 3 microns, and follows a linear relation of physical to genomic distance, consistent with an extended worm-like chain having a persistence length on the order of microns.

